# Plunge freezing tweezers with ultrasonic excitation for reducing preferred particle orientation in cryo-EM

**DOI:** 10.64898/2026.09.29.755300

**Authors:** Jakub Wenz, Mischa Flór, Constantin R. Krüger, Marcel Drabbels, Ulrich J. Lorenz

## Abstract

Preferred particle orientation, which results from the adsorption of proteins to the air-water interface, is a frequently encountered issue in single-particle cryo-electron microscopy (cryo-EM). Particles then present only a limited number of views, which are oversampled during data acquisition at the expense of throughput, while other views may be missing altogether. In severe cases, a high-resolution reconstruction becomes unattainable. We have recently established that ultrasonic excitation of the sample during vitrification improves the angular distribution by detaching particles from the air-water interface and reshuffling their orientations. Here, we describe an implementation of this principle for a plunge freezing apparatus, in which the ultrasonic transducer is integrated into the head of the pair of tweezers holding the sample, and show that this design is able to reduce preferred orientation. We demonstrate this for hemagglutinin, the canonical example of preferred orientation, and show that the fraction of side and tilted views increases with the excitation amplitude. This allows us to obtain a complete reconstruction of the trimer at 2.5 Å.

## Introduction

During the preparation of a cryo-EM specimen, many proteins adsorb to the air-water interface of the thin liquid film with hydrophobic patches on their surface.^1–3^ Vitrification then results in an angular distribution dominated by a small number of orientations, with others sparsely populated or absent entirely.^4^ This can limit the resolution along some viewing directions, and a proportionally larger number of micrographs has to be acquired to obtain an isotropic map, increasing cost and limiting the throughput of the technique. In the most severe cases, an interpretable map cannot be obtained at all.^5–8^

A variety of strategies have been devised to counter preferred orientation, but none has emerged as a general remedy. Collecting data with the stage tilted supplies missing views, but incurs the penalty that the electron beam has to take a longer path through the ice, which results in a poorer signal-to-noise ratio, which is a particular disadvantage for small proteins.^6,8–10^ A second family of approaches attempt to prevent adsorption to the interface^4,11–13^, either by adding surface-active compounds, by encapsulating the protein of interest in a protein shell^14,15^, or by using alternative specimen support film.^11,16–21^ However, these strategies require re-optimization of the sample preparation, which is laborious and may not be successful. A third route shortens the interval between sample application and vitrification so that fewer molecules have enough time to reach the interface.^22–28^ However, the instruments involved are frequently specialized and usually cannot outrun adsorption entirely. In a related approach, we have shown that flash melting a sample with a microsecond laser pulse and revitrifying it reduces preferred orientation.^29,30^ Our observations suggest that the laser pulse induces mechanical oscillations in the specimen support that detach particles from the interface through the accompanying motions of the liquid film. However, readsorption competes with this process, which is particularly fast for small proteins, limiting the method to larger particles.

Recently, we have shown that ultrasonic excitation provides a simpler approach for reducing preferred orientation by inducing mechanical oscillations in the sample (Fig. 1a).^31^ The specimen is continuously excited by an ultrasonic transducer, and the sample is vitrified with a jet of liquid ethane, which improves the angular distribution for proteins of a wide range of sizes. Electron tomography reveals that ultrasonic excitation detaches particles from the air-water interface, allowing them to reshuffle their orientations. Our experiments suggest that under continuous excitation, fluid motions induced by the oscillating specimen support establish a dynamic equilibrium of particle detachment and readsorption in which the particles populate orientations that were previously disfavored. We have also described an implementation of this scheme for a plunge freezing instrument, in which the specimen passes a series of ultrasonic transducers during plunging.^31^ In related work, others have shown that acoustic excitation with piezoelectric transducers glued to plunge freezing tweezers increases the particle density in the holes of the specimen support and distributes particles more evenly throughout the thickness of the ice.^32^ However, the effect on preferred orientation was not investigated. Here, we present an implementation in which a transducer is integrated into the head of the plunge freezing tweezers and show that it can overcome preferred orientation. Our implementation does not require any irreversible modification of the plunge freezing instrument, with the cable that powers the transducer routed over a pulley system that can be easily installed and removed. We demonstrate the approach on hemagglutinin, a canonical case of preferred orientation,^8,24,30,31,33–36^ and show that the angular distribution improves with increasing excitation amplitude.

**Figure 1.**
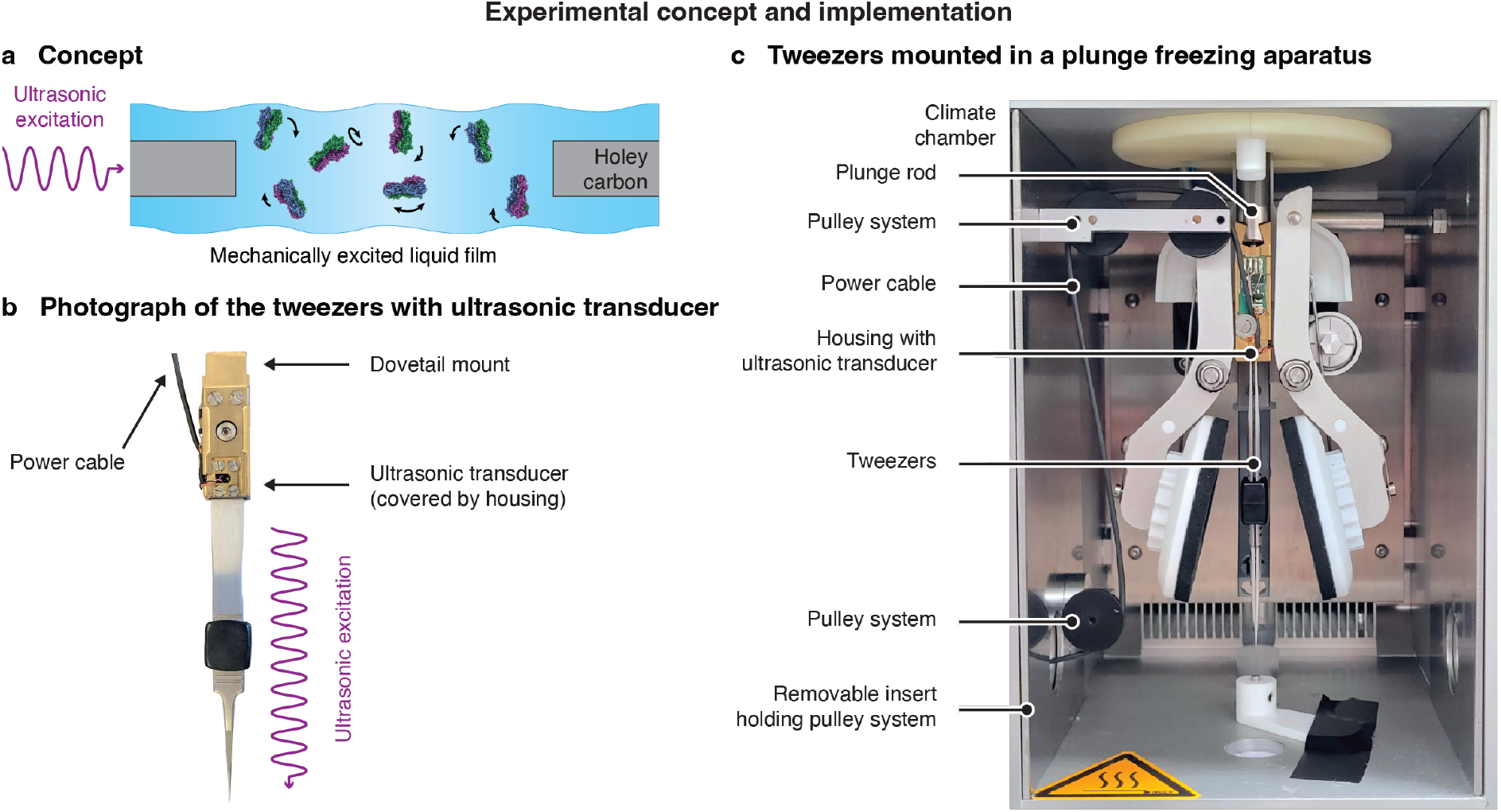
| Experimental concept and implementation. **a** Concept. Ultrasonic excitation of the sample induces motions of the liquid film that detach particles from the air-water interface, allowing them to reshuffle their orientations before the sample is vitrified. **b** Photograph of the modified tweezers. An ultrasonic transducer is integrated into the dovetail mount of the tweezers with which they are attached to the plunge rod of the plunge freezing instrument. The ultrasonic waves are transferred through the body of the tweezers to the sample. **c** Photograph of the tweezers mounted in the plunge freezing apparatus (Thermo Fisher Vitrobot Mark IV). The electrical cable required to power the ultrasonic transducer is guided by a pulley system that is mounted on a removable insert installed in the climate chamber.

## Results

A photograph of the modified tweezers is shown in Fig. 1b (Supplementary Information 1). The ultrasonic transducer (SECO SC049, driven by a 490 kHz square wave at either 10 V, 20 V, or 150 V amplitude) is integrated into a custom dovetail mount, machined from brass, with which the stainless-steel tweezers (Dumont L5 inox 08) are attached to the plunge rod of the plunge freezing instrument. Figure 1c shows a photograph of the tweezers mounted in a Thermo Fisher Vitrobot Mark IV. The cable required to power the tweezers is guided by a pulley system, which prevents it from getting tangled during plunging. The three wheels of the pulley system are mounted on two vertical aluminum boards, which are inserted on either side of the climate chamber of the plunge freezing device. A threaded rod, visible behind the blotting arms and plunge rod, pushes the boards against either wall of the chamber. The aluminum boards have circular holes that line up with the ports of the climate chamber through which the user can insert a pipette to apply the sample to the specimen grid. The left port is used to route the power cable out of the climate chamber. Mounting the pulley system in this fashion makes it possible to easily install and remove it, without any irreversible modifications of the plunge freezing apparatus.

We start the ultrasonic excitation of the sample during the plunging process, about 80 ms before the sample reaches the cryogen. As detailed in Supplementary Information 1, the transducer is triggered by a roller-lever microswitch that we have mounted in the housing of the plunge freezing instrument and that is actuated by the movement of the plunge rod. The transducer then operates continuously for 1 s before it switches off. Note that it is also possible to activate the ultrasonic transducer manually for several seconds before plunging the sample. In this case, a modification of the plunge freezing apparatus to install a trigger switch can be avoided.

We characterize the performance of the plunge freezing tweezers with integrated ultrasonic excitation for hemagglutinin, which frequently serves as a test case for methods to overcome preferred orientation. ^8,24,30,31,33–36^ As shown in Fig. 2a (top), a conventional sample plunge frozen on holey carbon support film (Quantifoil R1.2/1.3 on 200 mesh copper) has such strong preferred orientation, with top and bottom views dominating, that it only yields a truncated reconstruction with streaky features (inset) and a nominal resolution of 2.9 Å (Supplementary Information 2,3). To obtain the reconstruction, we use a well-established processing trick and perform the initial *ab initio* reconstruction only with particles from 2D class averages that correspond to side views as well as tilted views (Fig. 2b). However, these account for only 3 % and 8 % of particles, respectively. Accordingly, the map in Fig. 2a yields a low cFAR value of 0.02 and a low SCF* value of 0.32. Both metrics are used to assess the degree of preferred orientation and take values between 0 and 1, with 1 representing a perfectly isotropic distribution. The SCF* is based on the particle poses,^37^ while the cFAR uses a conical Fourier Shell Correlation to assess the anisotropy of the resulting map.^38^ Note that the angular distribution of the particles contained in the reconstruction provides only an unreliable estimate of the angular distribution of all particles in the sample since the volume against which the particles are aligned is truncated and since the processing workflow rejects many top and bottom views (Supplementary Information 3). We therefore estimate the angular distribution of the sample by merging the particles retained after 2D classification from all four samples in Fig. 2 and jointly refining them against a complete volume of hemagglutinin (Supplementary Information 4). As shown in Fig. 2c, we obtain an angular distribution for the conventional sample that is dominated by top and bottom views, with a low SCF* value of 0.40.

**Figure 2.**
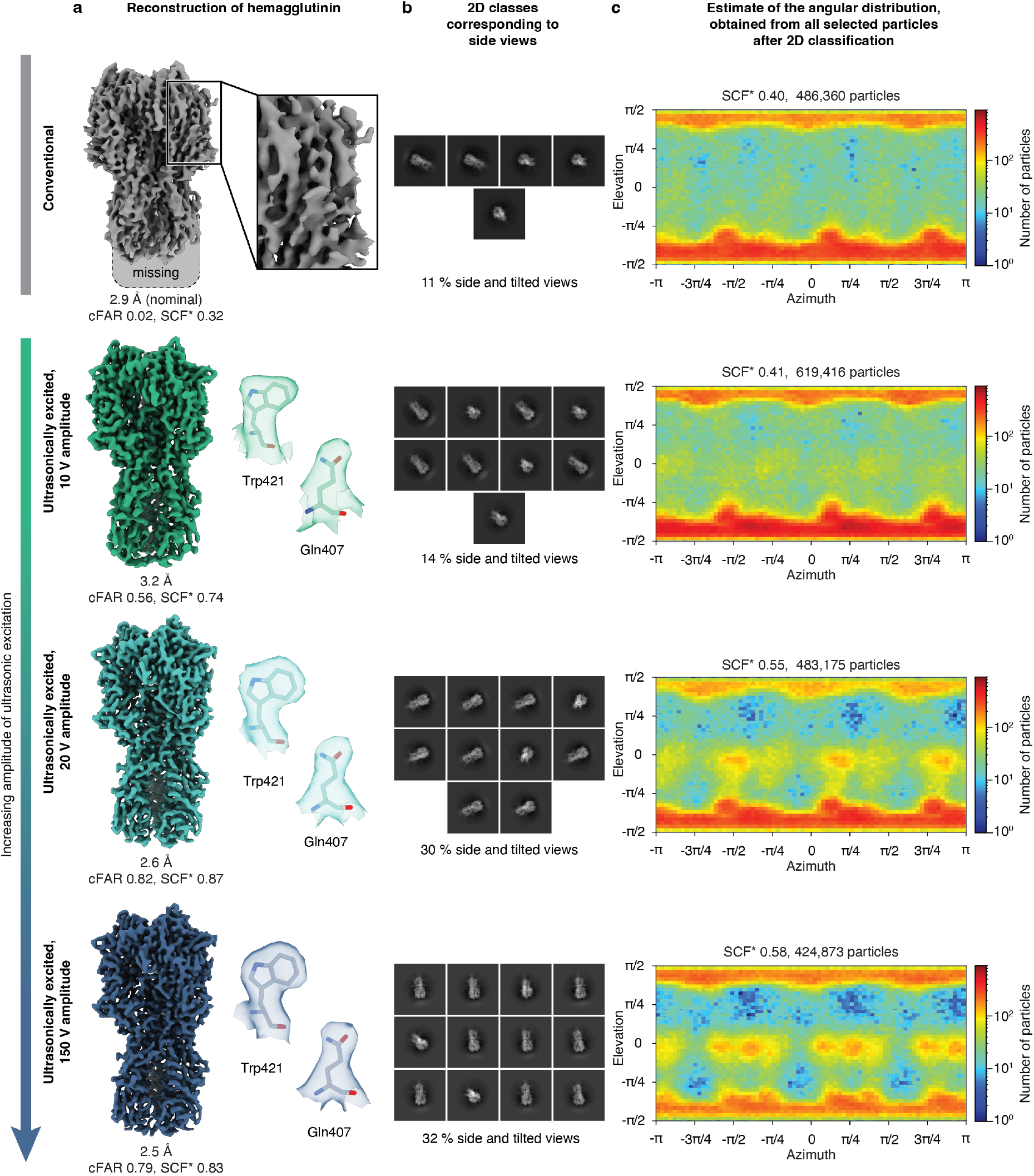
| The angular distribution of hemagglutinin improves with increasing amplitude of the ultrasonic excitation. **a** Reconstructions of hemagglutinin from a conventional sample as well as samples plunge frozen with ultrasonic excitation at amplitudes of 10 V, 20 V, and 150 V. While only a truncated map can be obtained from a conventional sample, ultrasonic excitation results in complete volumes. Details of the reconstructions are shown with a model (PDB ID 7VDF^34^) jiggle-fitted into the map. The map resolution is indicated together with cFAR and SCF* values. **b** Class averages corresponding to side and tilted views for each dataset. Their fraction increases from 11 % in a conventional sample to 30 % at high excitation amplitudes. **c** Estimates of the angular distribution in each sample. The estimates are obtained by merging the particles from all four samples that were retained after 2D classification and jointly refining them against the volume obtained with ultrasonic excitation at 20 V amplitude (Supplementary Information 4).

Ultrasonic excitation of the sample during plunge freezing improves the angular distribution of the particles and yields a complete map, in agreement with our previous experiments.^31^ At an excitation amplitude of 10 V (second row in Fig. 2), more side views are populated. Accordingly, the fraction of side and tilted views in the 2D class averages increases to 14 %, and a series of faint maxima appear in the angular distribution at an elevation of zero (the estimated SCF* value of the sample remains nearly unchanged at 0.41, while the SCF* value of the map increases to 0.74). This enables us to obtain a 3.2 Å reconstruction that is not truncated, with the cFAR significantly improved, to a value of 0.56. Selected side chain densities are shown.

The angular distribution of the particles improves further when we increase the amplitude of ultrasonic excitation. At an excitation amplitude of 20 V (third row in Fig. 2), the fraction of side and tilted views in the 2D class averages increases to 30 %, and the SCF* value of the sample improves to 0.55 (0.87 for the map). The resolution of the reconstruction accordingly improves to 2.6 Å, with the cFAR value increased further to 0.82. Raising the excitation amplitude to 150 V does not lead to a further significant improvement in the orientation distribution (fourth row in Fig. 2). The 2D class averages contain 32 % side and tilted views, and the SCF* value of the sample is 0.58 (0.83 for the map). The resulting 2.5 Å reconstruction has a cFAR value of 0.79.

As detailed in Supplementary Information 5, we obtain a similar result for the C-reactive protein. Ultrasonic excitation at 10 V amplitude generates a more even angular distribution and improves the resolution of the reconstruction from 3.3 Å for a conventional sample to 2.8 Å. If we increase the excitation amplitude to 150 V, the angular distribution changes, with different sets of side views now being preferentially populated. However, this does not result in a further improvement of the map resolution. For the 50S ribosomal subunit, ultrasonic excitation at 10 V or 150 V amplitude alters the orientation distribution, transferring population from the global maximum to several local maxima. However, the orientation statistics and map resolution remain similar, in contrast to our previous jet vitrification experiments, where ultrasonic excitation led to an improvement.^31^ This may be because in the present experiment, the conventional sample exhibits less orientation bias to begin with, comparable to the level achieved with ultrasonic excitation in the jet vitrification experiment.

Our experiments also provide further insights into the mechanism by which ultrasonic excitation alters the angular distribution. We have previously suggested that the increase in concentration of particles not attached to the interface must result from an increased rate at which particles detach or a decreased rate at which they reattach.^23,24,30^ Moreover, the relative change in attachment and detachment rates must differ between different particle orientations. For example, the increased population of hemagglutinin particles adsorbed in side views suggest that for these orientations, the increase in detachment rate or the decrease in attachment rate is smaller than for the top and bottom views. Our experiments add another detail to this picture by showing that this effect also depends on the excitation amplitude. For example, for the C-reactive protein, different sets of side views are preferentially populated at different amplitudes, while both appear evenly populated without ultrasonic excitation (Fig. S7). This suggests a complex mechanism by which mechanical excitation alters the rates of particle attachment and detachment.

In summary, we have presented a simple implementation of plunge freezing tweezers with ultrasonic excitation and have demonstrated their operation, notably showing that they reduce preferred orientation for hemagglutinin, a canonical case of preferred orientation, for which ultrasonic excitation yields a complete reconstruction at 2.5 Å resolution, whereas a conventional sample gives only a truncated map. Our implementation does not require any irreversible modification of the plunge freezing apparatus that might void its warranty, with the cable powering the ultrasonic transducer routed through a pulley system that is simply inserted into the climate chamber. We are currently working on dispensing with the power cable entirely and implementing a battery powered version of the tweezers to enable the routine and potentially widespread use of ultrasonic excitation during plunge freezing. It will only become clear over time which range of proteins can benefit from ultrasonic excitation. However, even if the angular distribution could only be improved for a small fraction of proteins, this would amount to important gains in throughput, with the largest benefits for difficult cases such as hemagglutinin. Also note that ultrasonic excitation can be easily used together with other methods for reducing preferred orientation, potentially combining the effects. In our experiments, we obtain no significant improvement by increasing the excitation amplitude beyond 20 V. Future experiments will have to show if this holds for a wide range of proteins.

## Author contributions

**Jakub Wenz:** Conceptualization (equal), Investigation (lead), Formal analysis (lead), Data curation (equal), Writing – original draft (equal), Writing – Review & Editing (equal), Visualization (lead). **Mischa Flór:** Methodology (lead), Investigation (supporting), Writing – Review & Editing (equal), Visualization (supporting), Project administration (equal), Funding acquisition (equal). **Constantin R. Krüger:** Methodology (supporting), Writing – Review & Editing (equal), Project administration (equal), Funding acquisition (equal). **Marcel Drabbels:** Conceptualization (equal), Data curation (equal), Writing – Review & Editing (equal), Supervision (equal), Project administration (equal). **Ulrich J. Lorenz:** Conceptualization (equal), Writing – original draft (equal), Writing – Review & Editing (equal), Supervision (equal), Project administration (equal), Funding acquisition (equal).

## Data and materials availability

Supplementary Information is available for this paper.

The cryo-EM maps have been deposited in the Electron Microscopy Data Bank (EMDB) and the Electron Microscopy Public Image Archive (EMPIAR) under accession codes EMD-100126 and EMPIAR-XXXXXX (recombinant hemagglutinin trimer; conventional sample), EMD-100127 and EMPIAR-XXXXXX (recombinant hemagglutinin trimer; ultrasonically excited sample, 10 V amplitude), EMD-100133 and EMPIAR-XXXXX (recombinant hemagglutinin trimer; ultrasonically excited sample, 20 V amplitude), EMD-100134 and EMPIAR-XXXXX (recombinant hemagglutinin trimer; ultrasonically excited sample, 150 V amplitude), EMD-100107 and EMPIAR-XXXXXX (human C-reactive protein; conventional sample), EMD-100124 and EMPIAR-XXXXXX (human C-reactive protein; ultrasonically excited sample, 10 V amplitude), EMD-100125 and EMPIAR-XXXXXX (human C-reactive protein; ultrasonically excited sample, 150 V amplitude), EMD-100096 and EMPIAR-XXXXXX (50S ribosomal subunit; conventional sample), EMD-100097 and EMPIAR-XXXXXX (50S ribosomal subunit; ultrasonically excited sample, 10 V amplitude), and EMD-100108 and EMPIAR-XXXXXX (50S ribosomal subunit; ultrasonically excited sample, 150 V amplitude).

## Acknowledgments

The authors would like to thank the EPFL ISIC electrical and mechanical workshops for their help in realizing the project, in particular S. Dutoit, B.C. Le Geyt, and G. Pasche, as well as B. Beckert from the Dubochet Center for Imaging Lausanne for providing a sample of the 50S ribosomal subunit. Cryo-EM data collection was performed at the Dubochet Center for Imaging Lausanne (a joint initiative from EPFL, UNIGE, UNIL, UNIBE) with the assistance of B. Beckert, S. Nazarov, I. Mohammed, and E. Uchikawa. The work was supported by Swiss National Science Foundation Grants TMCG-2_213773 (UJL) and Innosuisse Bridge PoC 40B1-0_238531 (CRK), and EPFL Innogrant (CRK, MF). Use of artificial intelligence (AI): Claude (Anthropic) and Perplexity (Perplexity AI) were used for literature search and language editing. All literature sources obtained with AI assistance were independently checked by the authors against the original publications. The authors reviewed and approved all AI-assisted edits of the text.

