## Supplementary information for "Plunge freezing tweezers with ultrasonic excitation for reducing preferred particle orientation in cryo-EM"

This Supplementary Information contains:

1 | Timing of ultrasonic excitation

2 | Cryo-EM sample preparation

3 | Data collection and processing – hemagglutinin

4 | Estimate of the angular distribution in the conventional and ultrasonically excited hemagglutinin samples

5 | Plunge freezing with ultrasonic excitation for samples of the C-reactive protein and the 50S ribosomal subunit

6 | Data collection and processing – C-reactive protein

7 | Data collection and processing – 50S ribosomal subunit

8 | Supplementary tables

### 1 | Timing of ultrasonic excitation

The ultrasonic excitation of the sample is started during the plunging process with the help of the roller-lever switch, which is actuated by the movement of the plunge rod. As shown in the photograph of Fig. S1, the switch is mounted on the back panel of the plunge freezing apparatus, near the vertical rail on which the plunge rod travels. The motion of the plunge rod is controlled by a stepper motor, which drives a belt that is attached to the topmost linear bearing on which the plunge rod is mounted. As the plunge rod descends, a white, plastic tab that we have glued to the fixture of the belt actuates the roller-lever switch, triggering the ultrasonic excitation of the sample for the duration of 1 s. From the placement of the switch and the speed with which the plunge rod moves, we estimate that the grid is excited for about 80 ms before it dips in to the cryogen.

**Triggering of ultrasonic excitation during plunging**

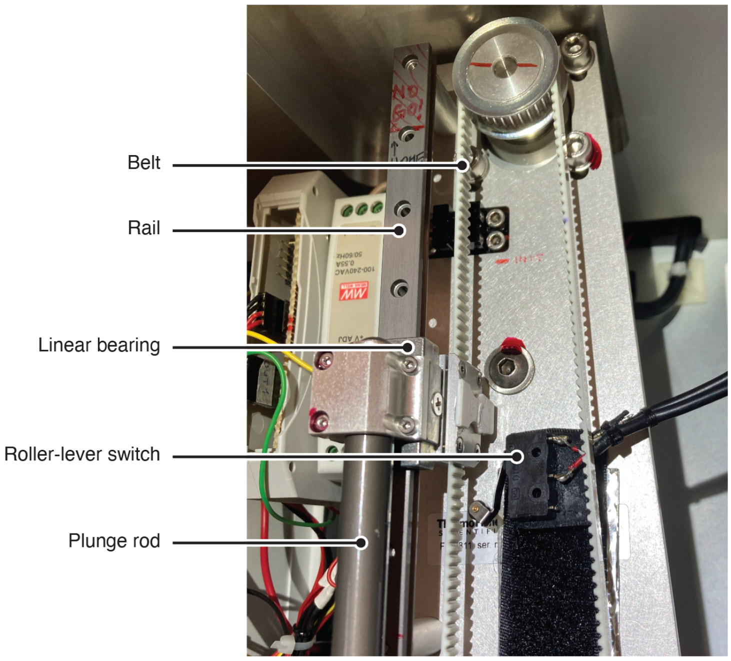


**Figure S1 | Photograph of the roller-lever switch that triggers ultrasonic excitation during the plunging process.** The plunge rod travels on a linear rail and is moved by a belt, driven by a stepper motor. As the plunge rod descends, a roller-lever switch is actuated to start ultrasonic excitation. The switch is mounted on the back panel of the plunge freezing apparatus and is activated by a white, plastic tab that is glued onto the fixture that connects the belt to the topmost linear bearing on which the rod is mounted.

**2 | Cryo-EM sample preparation**

Purified hemagglutinin trimer recombinant protein (1 mg/mL in 1x PBS, pH 7.5) was purchased from MyBioSource (catalogue number MBS434205) and used as is. C-reactive protein from human fluids was purchased from Sigma-Aldrich and used as is (C4603; 2.6 mg/mL, 20 mM Tris buffer, pH 7.8–8.2, 280 mM NaCl, 5 mM $CaCl_2$, 0.1 % $NaN_3$). A purified sample of the 50S ribosomal subunit (40 $OD_{260}$/mL, 20 mM HEPES buffer, pH 7.5, 100 mM NaCl, 2 mM $MgCl_2$) was kindly provided to us by Dr. Bertrand Beckert from the Dubochet Center for Imaging in Lausanne.

Cryo-EM samples were prepared by applying 3.5 µL of the protein solution onto holey carbon grids (R1.2/1.3, 200 Cu mesh, Quantifoil) that were rendered hydrophilic using air glow discharge (90 s, 15 mA, EasiGlow, TedPella). Samples were prepared using a Thermo Fisher Vitrobot MkIV plunge-freezing device (4 °C, 90 % relative humidity, blotting force 10, 4.5 s blotting time for hemagglutinin; 20 °C, 95 % relative humidity, blotting force 10, 3.5 s blotting time for C-reactive protein; 20 °C, 95 % relative humidity, blotting force 0, 2 s blotting time for 50S ribosomal subunit) and vitrified in liquid ethane. All samples were perpared with the modified tweezers shown in Fig. 1b. For the conventional samples, the ultrasonic transducer was not activated during plunging.

**3 | Data processing and analysis – hemagglutinin**

High-resolution micrographs of all samples of hemagglutinin trimer (conventional and ultrasonically excited) were collected at the Dubochet Center for Imaging in Lausanne, Switzerland, using a Thermo Fisher Titan Krios G4i transmission electron microscope equipped with a Falcon IV direct electron detector. Single particle reconstructions were performed using CryoSPARC v5.0.6[1] with the corresponding workflows illustrated in Figs. S2–S5. The data acquisition parameters and processing statistics are summarized in Supplementary table 1. Visualization of the density maps was performed using Chimera X[2,3].

Micrographs of all datasets were first subjected to patch motion correction and patch CTF estimation. Micrographs with estimated CTF fit worse than 10 Å were rejected. Particles from conventional and ultrasonically excited samples were picked using a blob picker (diameter 50 – 200 Å), extracted with a box size of 384 px and Fourier-cropped to 192 px.

For the conventional sample, the particles were first subjected to two rounds of 2D classification. After the final 2D classification, particles corresponding to all good classes were selected, from which particles corresponding to side and tilted views, as well as top/bottom views, were selected separately. The particle stack corresponding to side and tilted views was reconstructed using an *ab initio* with three classes ($C_1$ symmetry) followed by a heterogeneous refinement ($C_3$ symmetry). The particles belonging to the best class were non-uniformly refined ($C_3$ symmetry), producing a nearly complete low-resolution volume. Particles representing top/bottom views were reconstructed using an *ab* initio with a single class ($C_1$ symmetry), followed by a non-uniform refinement ($C_3$ symmetry), producing a truncated volume. All selected particles were subjected to two rounds of heterogeneous refinement ($C_3$ symmetry) using one full and one truncated volume and two decoy volumes as an input. Particles from the best class were then re-extracted using a full box size of 384 px and subjected to a non-uniform refinement ($C_3$ symmetry), followed by a further round of heterogeneous refinement ($C_3$ symmetry) using two copies of the volume from the previous non-uniform refinement and one truncated volume. Finally, particles belonging to the best class were non-uniformly refined ($C_3$ symmetry). The final volume from the conventional sample is shown in Fig. S2.

The dataset from the 10 V ultrasonically excited sample was processed as follows. First, four rounds of 2D classification were performed to clean the particle stack. From the last 2D classification, particles corresponding to all good classes were selected, from which particles corresponding to side and tilted views, as well as top/bottom views, were selected separately. The particle stack corresponding to side and tilted views was subjected to *ab initio* reconstruction with 3 classes ($C_1$ symmetry), followed by a heterogeneous refinement ($C_3$ symmetry). Particles belonging to the best class were non-uniformly refined ($C_3$ symmetry), producing a reconstruction of a full volume. Similarly, particles that represented top/bottom views were reconstructed using an *ab initio* (single class, $C_1$ symmetry) followed by a non-uniform refinement ($C_3$ symmetry) to produce the truncated volume similar to the conventional sample. All selected particles were then subjected to two rounds of heterogeneous refinement ($C_3$ symmetry) with a full volume (from side and tilted views), truncated volume (from top/bottom views) and two decoy classes as input. Particles belonging to the best class were subjected to two more rounds of heterogeneous refinement ($C_3$ symmetry) with two full and one truncated volume as inputs. The final particle stack reconstructing into a full volume, containing only 41,316 particles, was then re-extracted using a full box size of 384 px and subjected to final non-uniform refinement ($C_3$ symmetry), producing a volume displayed in Fig. S3.

The datasets of 20 V and 150 V ultrasonically excited samples were processed using the following identical protocol. Particles were initially subjected to three rounds of 2D classification. From the last 2D classification, particles corresponding to all good classes were selected, from which particles corresponding to side and tilted views, as well as top/bottom views, were selected separately. The particle stack corresponding to side and tilted views was reconstructed using an *ab initio* with three classes ($C_1$ symmetry) followed by a heterogeneous refinement ($C_3$ symmetry). The particles belonging to the best class were non-uniformly refined ($C_3$ symmetry), producing a full volume. Particles representing top/bottom views were reconstructed using an *ab* initio with a single class ($C_1$ symmetry), followed by a non-uniform refinement ($C_3$ symmetry), producing a truncated volume. All selected particles were subjected to two rounds of heterogeneous refinement ($C_3$ symmetry) using one full and one truncated volume and two decoy volumes as an input. Particles from the best class were then re-extracted using a full box size of 384 px and subjected to a non-uniform refinement ($C_3$ symmetry), followed by a further round of heterogeneous refinement ($C_3$ symmetry) using three copies of the volume obtained from the previous non-uniform refinement as an input. Finally, the particles belonging to the best class (around 80,000 particles for each dataset) were non-uniformly refined ($C_3$ symmetry). The final reconstructions are shown in Figs. S4 and S5 for samples ultrasonically excited with 20 V and 150 V amplitude, respectively.

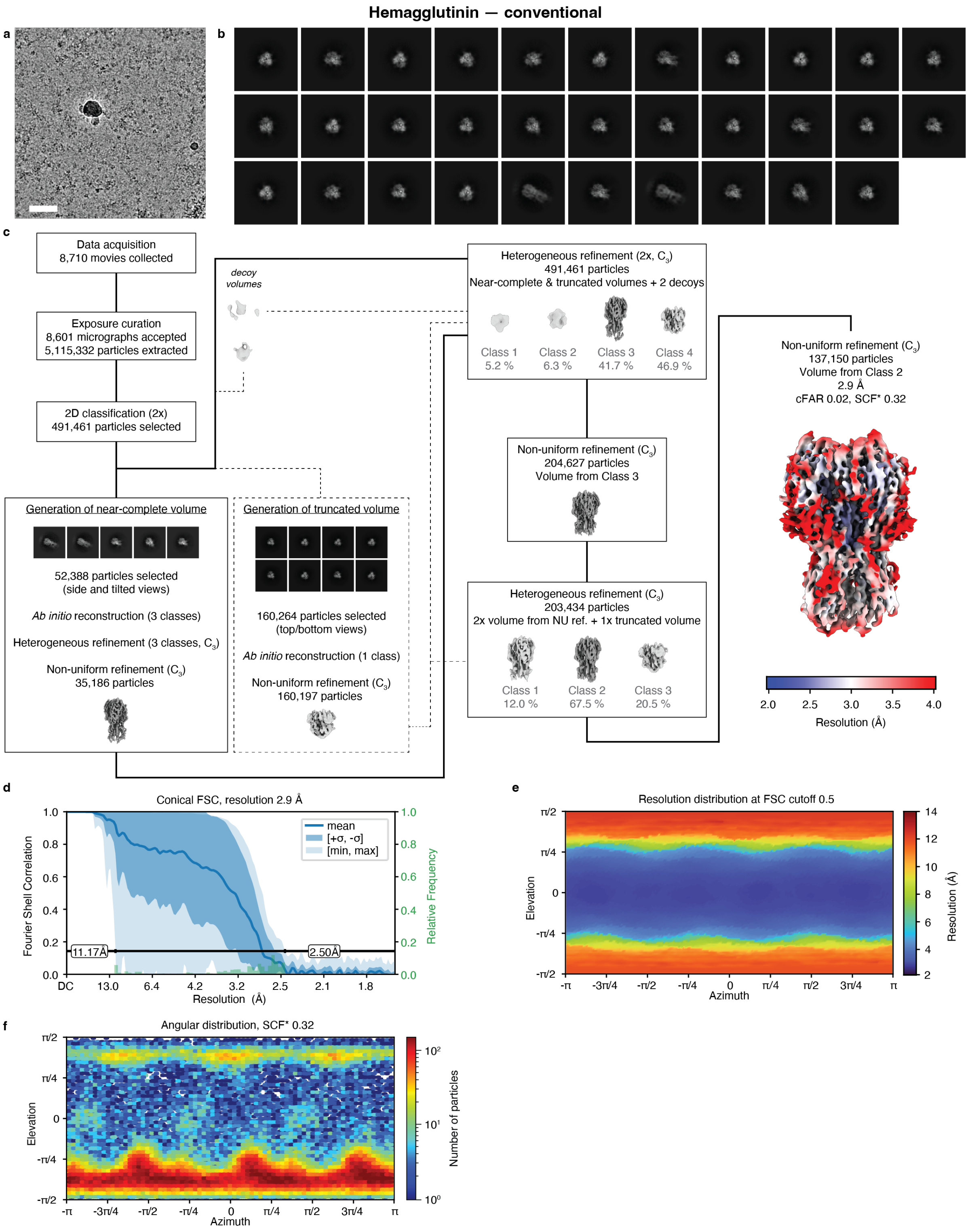


**Figure S2 | Processing workflow for the conventional sample of hemagglutinin. a** Representative micrograph. Scale bar, 500 Å. **b** Selected 2D class averages. **c** Data processing workflow in CryoSPARC. Symmetry applied is indicated in the parentheses. The volumes are displayed at a level of 10σ above the mean. The final map is displayed at a level 10σ above the mean, with the local resolution estimation at the FSC cutoff of 0.5 indicated in color. **d** Conical FSC, with the 0.143 threshold indicated by a black line. The mean FSC value is shown as a solid blue line. Dark blue shading indicates one standard deviation of the FSC curve, while the light blue shading represents the minimum and maximum FSC values. A histogram of the resolution values obtained from the conical FSC is shown in green. **e** Angular distribution of the resolution at FSC cutoff 0.5, generated using the 3DFSC file from Orientation Diagnostics job in CryoSPARC. **f** Angular distribution of the final reconstruction.

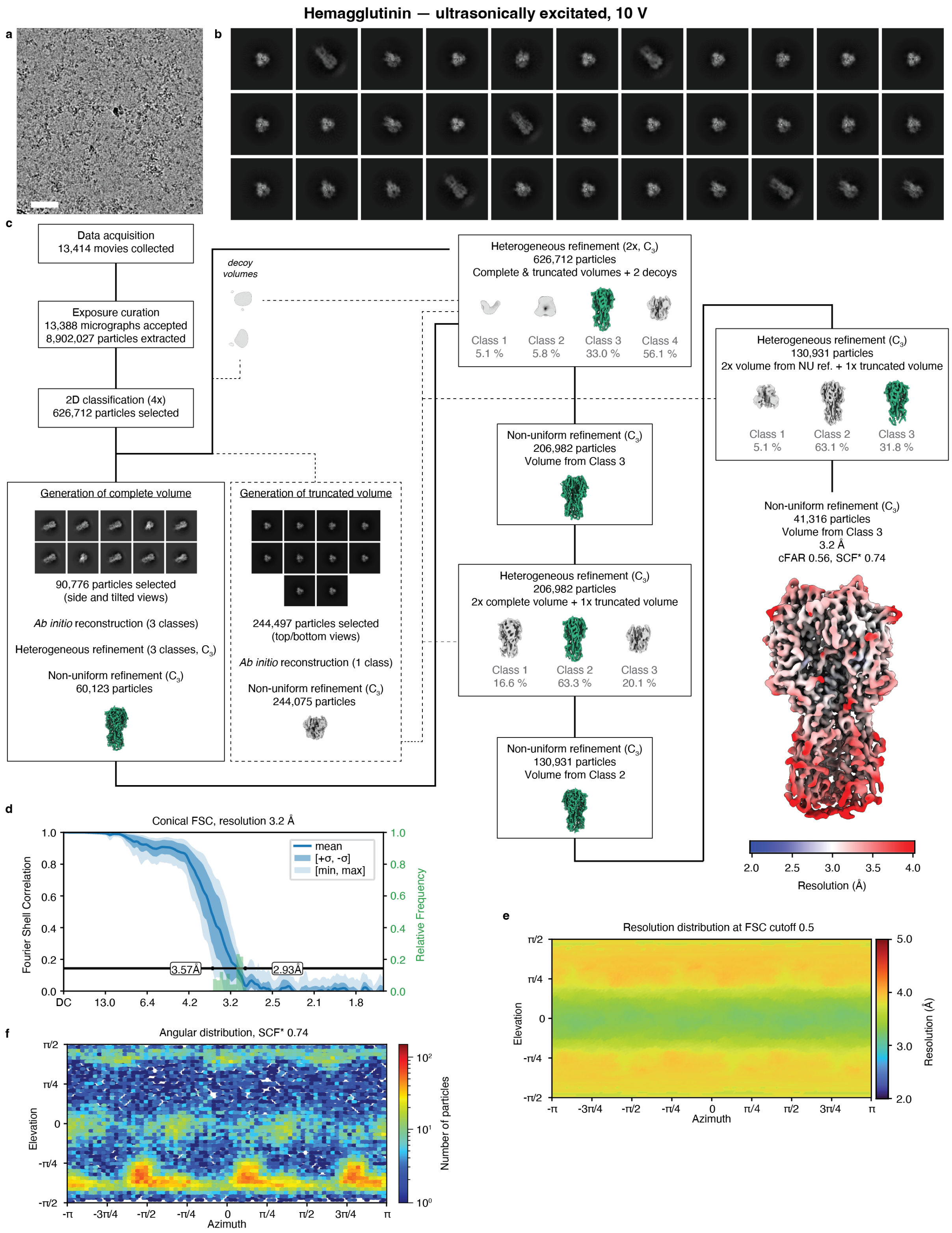


**Figure S3 | Processing workflow for the ultrasonically excited (10 V amplitude) sample of hemagglutinin. a** Representative micrograph. Scale bar, 500 Å. **b** Selected 2D class averages. **c** Data processing workflow in CryoSPARC. Symmetry applied is indicated in the parentheses. The volumes

are displayed at a level of 10$\sigma$ above the mean. The final map is displayed at a level 10$\sigma$ above the mean, with the local resolution estimation at the FSC cutoff of 0.5 indicated in color. **d** Conical FSC, with the 0.143 threshold indicated by a black line. The mean FSC value is shown as a solid blue line. Dark blue shading indicates one standard deviation of the FSC curve, while the light blue shading represents the minimum and maximum FSC values. A histogram of the resolution values obtained from the conical FSC is shown in green. **e** Angular distribution of the resolution at FSC cutoff 0.5, generated using the 3DFSC file from Orientation Diagnostics job in CryoSPARC. **f** Angular distribution of the final reconstruction.

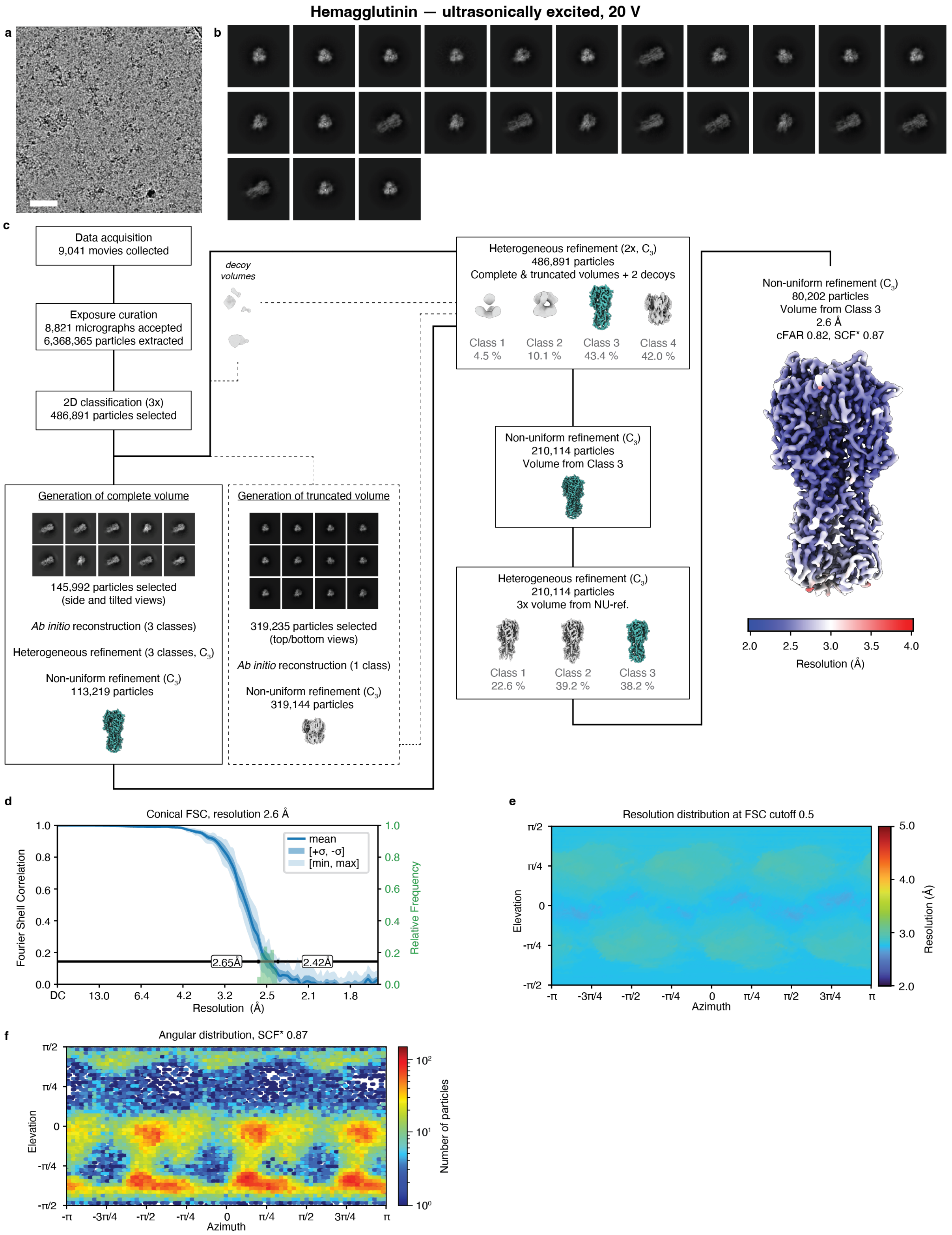


**Figure S4 | Processing workflow for the ultrasonically excited (20 V amplitude) sample of hemagglutinin. a** Representative micrograph. Scale bar, 500 Å. **b** Selected 2D class averages. **c** Data processing workflow in CryoSPARC. Symmetry applied is indicated in the parentheses. The volumes

are displayed at a level of 10$\sigma$ above the mean. The final map is displayed at a level 10$\sigma$ above the mean, with the local resolution estimation at the FSC cutoff of 0.5 indicated in color. **d** Conical FSC, with the 0.143 threshold indicated by a black line. The mean FSC value is shown as a solid blue line. Dark blue shading indicates one standard deviation of the FSC curve, while the light blue shading represents the minimum and maximum FSC values. A histogram of the resolution values obtained from the conical FSC is shown in green. **e** Angular distribution of the resolution at FSC cutoff 0.5, generated using the 3DFSC file from Orientation Diagnostics job in CryoSPARC. **f** Angular distribution of the final reconstruction.

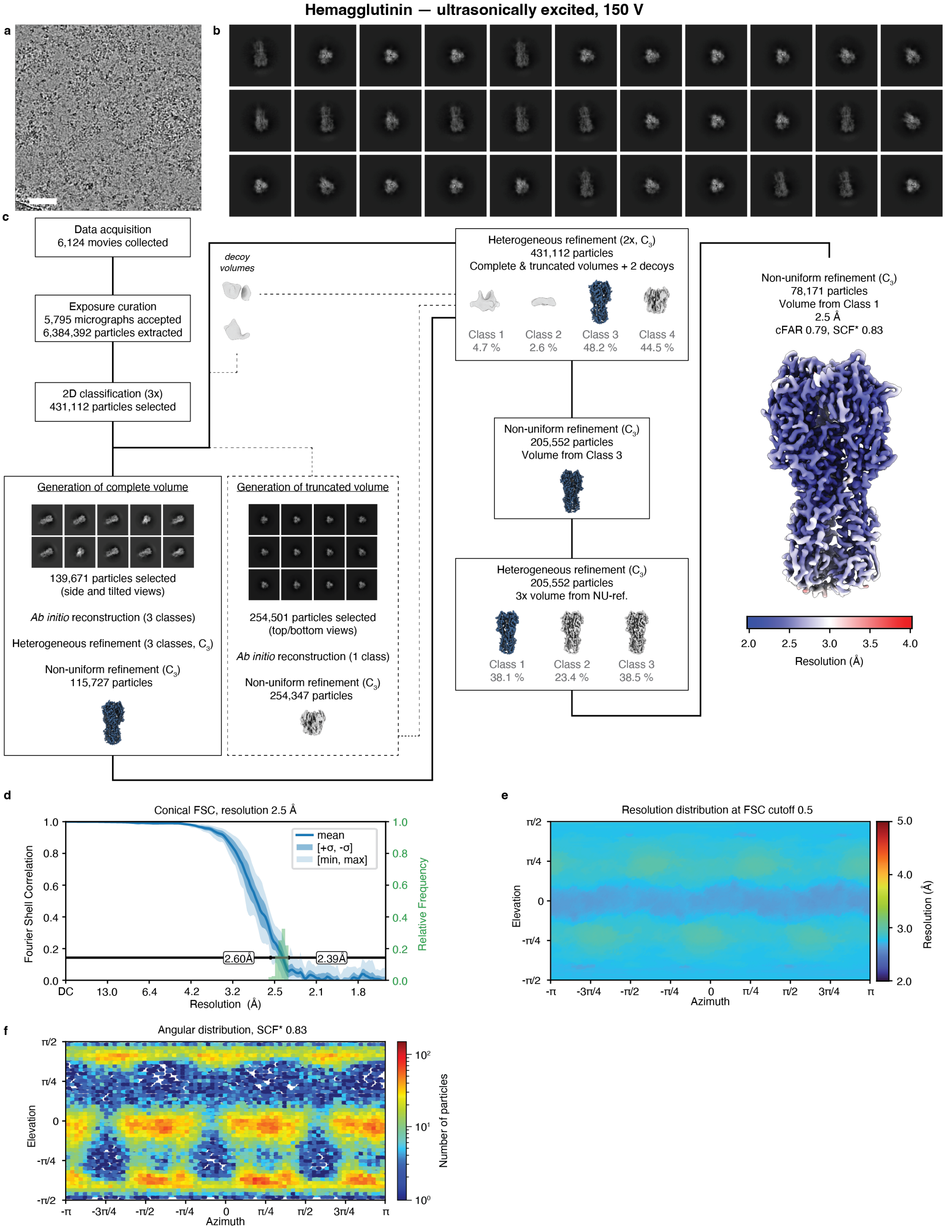


**Figure S5 | Processing workflow for the ultrasonically excited (150 V amplitude) sample of hemagglutinin.** **a** Representative micrograph. Scale bar, 500 Å. **b** Selected 2D class averages. **c** Data processing workflow in CryoSPARC. Symmetry applied is indicated in the parentheses. The volumes

are displayed at a level of 10$\sigma$ above the mean. The final map is displayed at a level 10$\sigma$ above the mean, with the local resolution estimation at the FSC cutoff of 0.5 indicated in color. **d** Conical FSC, with the 0.143 threshold indicated by a black line. The mean FSC value is shown as a solid blue line. Dark blue shading indicates one standard deviation of the FSC curve, while the light blue shading represents the minimum and maximum FSC values. A histogram of the resolution values obtained from the conical FSC is shown in green. **e** Angular distribution of the resolution at FSC cutoff 0.5, generated using the 3DFSC file from Orientation Diagnostics job in CryoSPARC. **f** Angular distribution of the final reconstruction.

**4 | Estimate of the angular distribution in the conventional and ultrasonically excited hemagglutinin samples**

Figure 2a shows the reconstructions of hemagglutinin obtained from a conventional sample as well as the ultrasonically excited samples. As detailed in Supplementary Information 3, the workflows yielding these reconstructions involve several rounds of heterogeneous refinement that remove many top and bottom views from the particle stack. The angular distributions of the remaining particles, which are shown in the workflows, therefore do not accurately reflect the angular distributions of the sample. To obtain a better estimate of these distributions, we merged the particles retained after 2D classification from all four datasets and jointly refined them against the complete volume of hemagglutinin obtained from the sample that was ultrasonically excited with 20 V amplitude. The corresponding workflow is illustrated in Fig. S6a, with the resulting estimates of the angular distributions in Fig. S6b (from Fig. 2c). Ultrasonic excitation visibly populates more side views, with the SCF* values increasing accordingly.

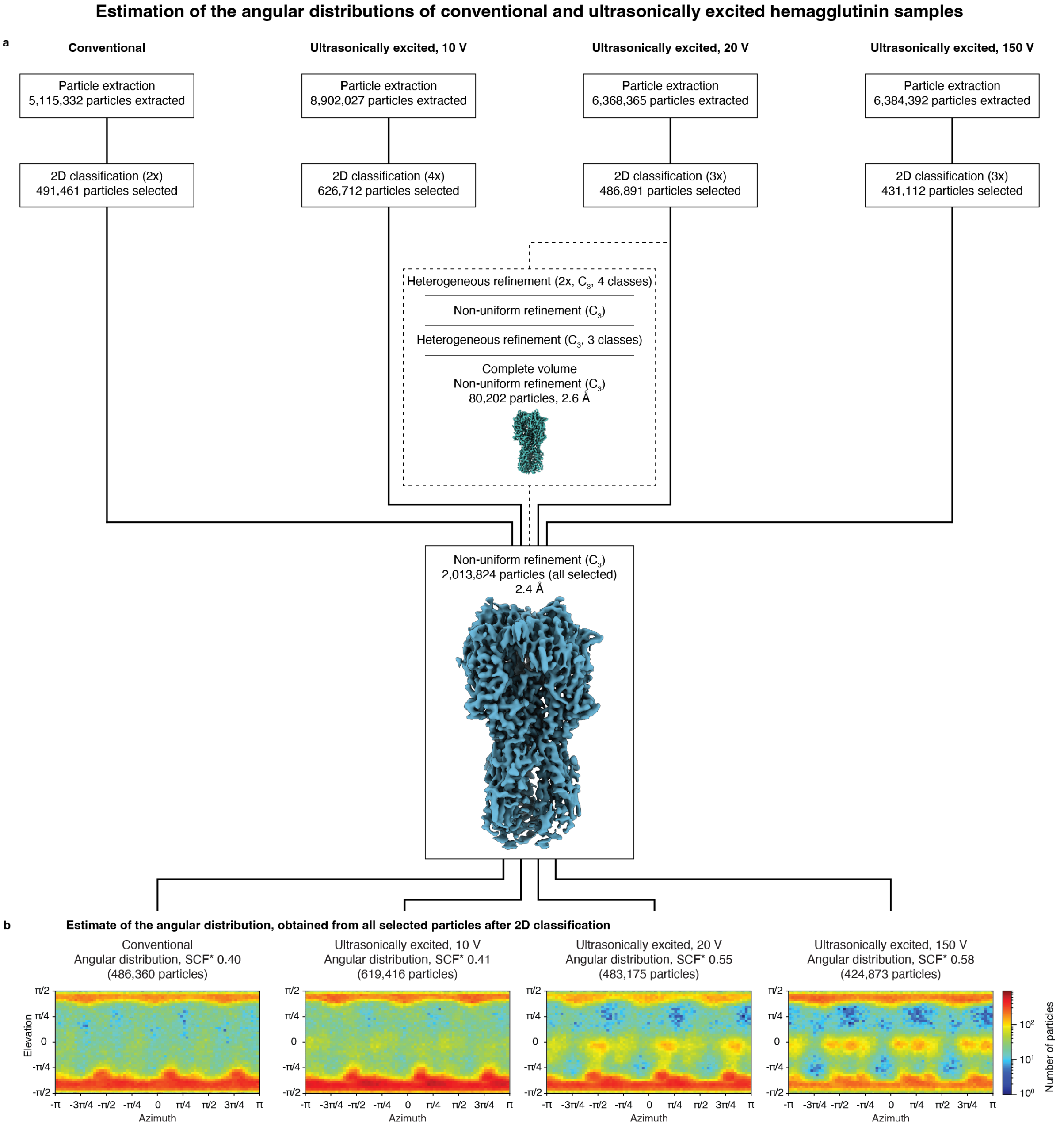


**Figure S6 | Estimate of the angular distributions in conventional and ultrasonically excited hemagglutinin samples. a** Data processing workflow in CryoSPARC. The particles retained after 2D classification from all four datasets are merged and jointly refined against the complete volume of obtained from the sample that was ultrasonically excited with 20 V amplitude. All maps are displayed at a level of 10σ above the mean. The symmetry applied is indicated in parentheses. **b** Resulting estimates of the angular distributions (as in Fig. 2c).

### 5 | Plunge freezing with ultrasonic excitation for samples of the C-reactive protein and the 50S ribosomal subunit

Ultrasonic excitation during plunge freezing similarly improves the angular distribution of C-reactive protein, in agreement with our previous experiments, where we employed jet vitrification.[4] Figure S7 shows reconstructions and angular distributions of a conventional sample together with samples plunged with ultrasonic excitation at 10 V (150 V) excitation amplitude (Supplementary Information 6). Ultrasonic excitation alters the angular distribution, changing the relative population between different side views as well as top and bottom views and improving the SCF* value from 0.82 to 0.96 (0.89). Accordingly, the resolution of the reconstruction improves from 3.3 Å to 2.8 Å (2.8 Å), with the cFAR value increasing from 0.62 to 0.78 (0.71). Interestingly, the angular distributions at 10 V and 150 V excitation amplitude show qualitative differences, with top and bottom views depopulated at 10 V compared to a conventional sample, while top views are prominently present at 150 V. Moreover, while both angular distributions feature an increased number of side views, different side views are populated at 10 V and at 150 V excitation amplitude.

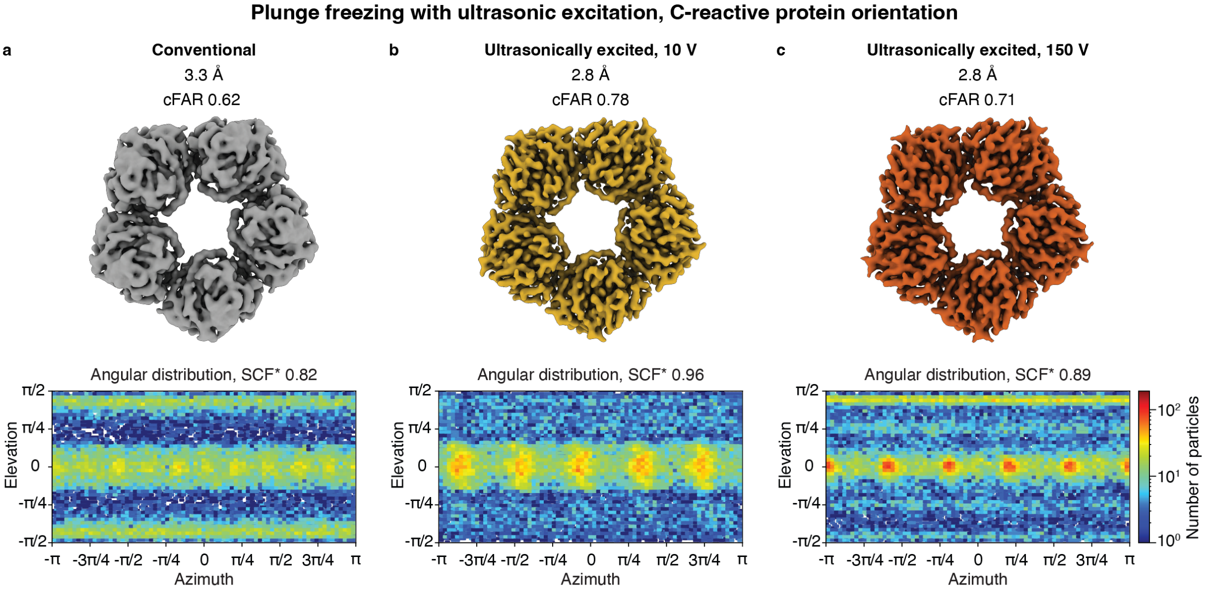


**Figure S7 | Plunge freezing with ultrasonic excitation, C-reactive protein. a** Reconstruction and angular distribution for a conventional sample. **b,c** Ultrasonic excitation with 10 V and 150 V excitation amplitude, respectively, improves the angular distribution, resulting in reconstructions with better resolution.

Ultrasonic excitation during plunge freezing alters the angular distribution of the 50S ribosomal subunit only slightly (Supplementary Information 7), increasing the population in some of the local maxima, without significantly changing the SCF* value or the resolution of the reconstruction (Fig. S8). The cFAR value increases slightly from 0.63 in a conventional sample to 0.71 (0.67) at 10 V (150 V) excitation amplitude.

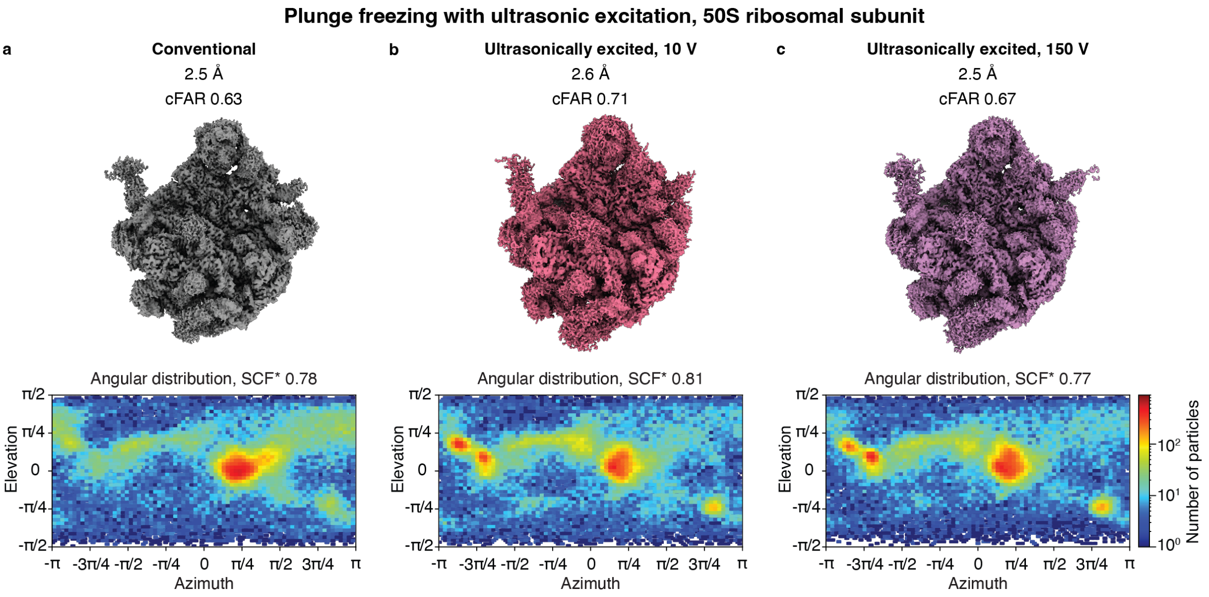


**Figure S8 | Plunge freezing with ultrasonic excitation, 50S ribosomal subunit. a** Reconstruction and angular distribution for a conventional sample. **b,c** Ultrasonic excitation with 10 V and 150 V excitation amplitude, respectively, slightly alters the angular distribution, but does not lead to an averall improvement.

**6 | Data processing and analysis – C-reactive protein**

High-resolution micrographs of all samples of C-reactive protein (conventional and ultrasonically excited) were collected at the Dubochet Center for Imaging in Lausanne, Switzerland, using a Thermo Fisher Titan Krios G4i transmission electron microscope equipped with a Falcon IV direct electron detector. Single particle reconstructions were performed using CryoSPARC v5.0.6[1] with the corresponding workflows illustrated in Figs. S9–S11. The data acquisition parameters and processing statistics are summarized in Supplementary table 2. Visualization of the resulting density maps was performed using Chimera X[2,3].

Micrographs were first subjected to patch motion correction and patch CTF estimation. Micrographs with estimated CTF fit worse than 8 Å were rejected. Particles from conventional and ultrasonically excited samples were picked using a blob picker (diameter 50 – 150 Å), extracted with a box size of 384 px and Fourier-cropped to 192 px.

All datasets were processed using the following workflow. Three rounds (conventional and 10 V ultrasonically excited samples) or five rounds (150 V ultrasonically excited sample) of 2D classification were first used to clear the particle stack with only pentamers left for further processing. Decameric particles were removed during the 2D classification and not processed further, as those do not suffer from preferred orientation and therefore are not the target of this study. The selected particles were subjected to *ab initio* reconstruction with three classes, followed by a homogeneous refinement ($C_5$ symmetry) of those classes. Subsequently, two rounds of heterogeneous refinement were performed with the selected particles and three volumes originating from the previous homogeneous refinement used as an input and $C_5$ symmetry imposed. Finally, a set of 50,000 particles belonging to the best class was selected randomly and non-uniformly refined to produce the final reconstruction shown in Fig. S9 (conventional) and Figs. S10 and S11 (samples ultrasonically excited with 10 V and 150 V amplitude, respectively).

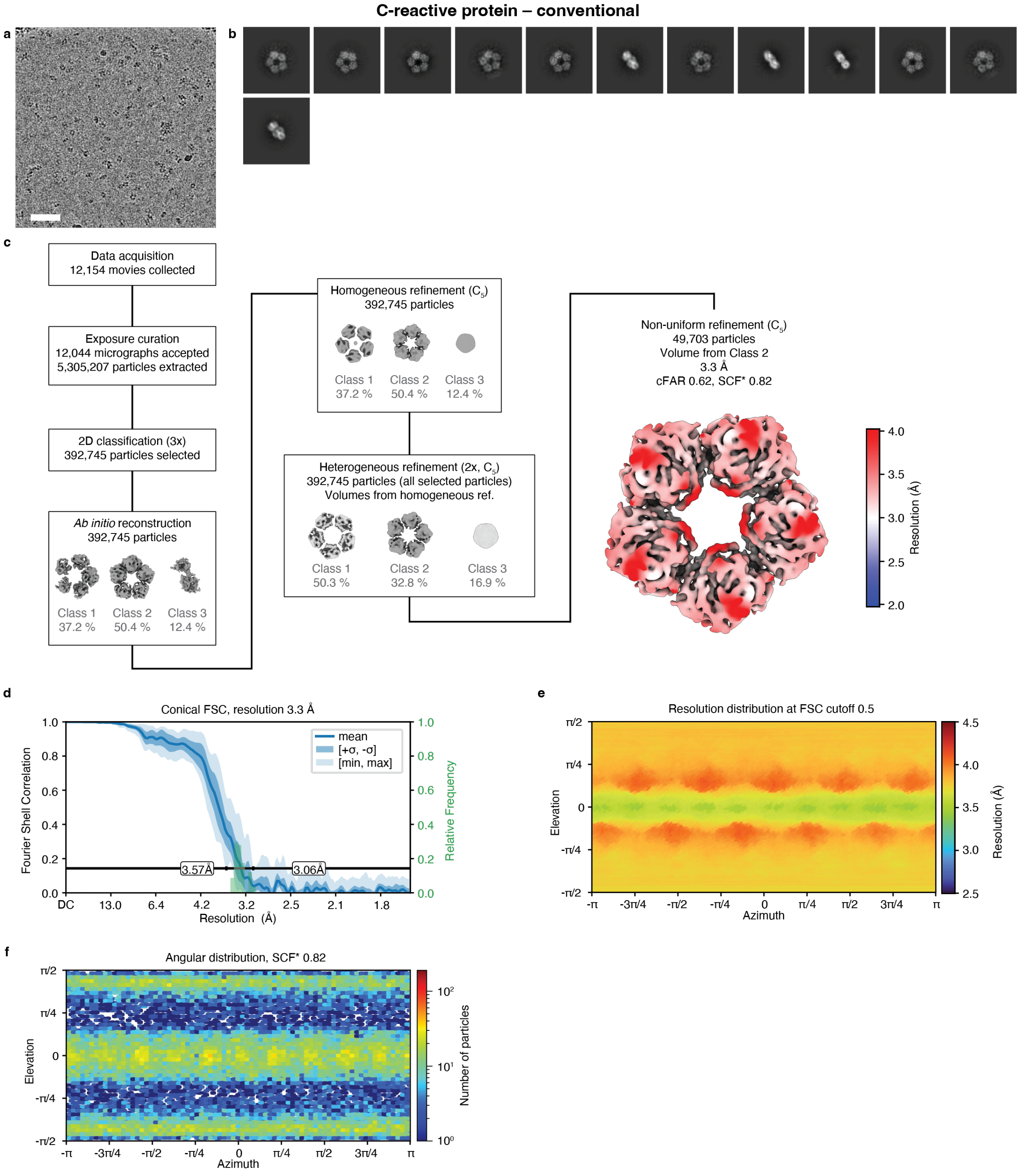


**Figure S9 | Processing workflow for the conventional sample of C-reactive protein.** **a** Representative micrograph. Scale bar, 500 Å. **b** Selected 2D class averages. **c** Data processing workflow in CryoSPARC. Symmetry applied is indicated in the parentheses. The volumes are displayed at a level of 10σ above the mean. The final map is displayed at a level 10σ above the mean, with the local resolution estimation at the FSC cutoff of 0.5 indicated in color. **d** Conical FSC, with the 0.143 threshold indicated by a black line. The mean FSC value is shown as a solid blue line. Dark blue shading indicates one standard deviation of the FSC curve, while the light blue shading represents the minimum and maximum FSC values. A histogram of the resolution values obtained from the conical FSC is shown in green. **e** Angular distribution of the resolution at FSC cutoff 0.5, generated using the 3DFSC file from Orientation Diagnostics job in CryoSPARC.

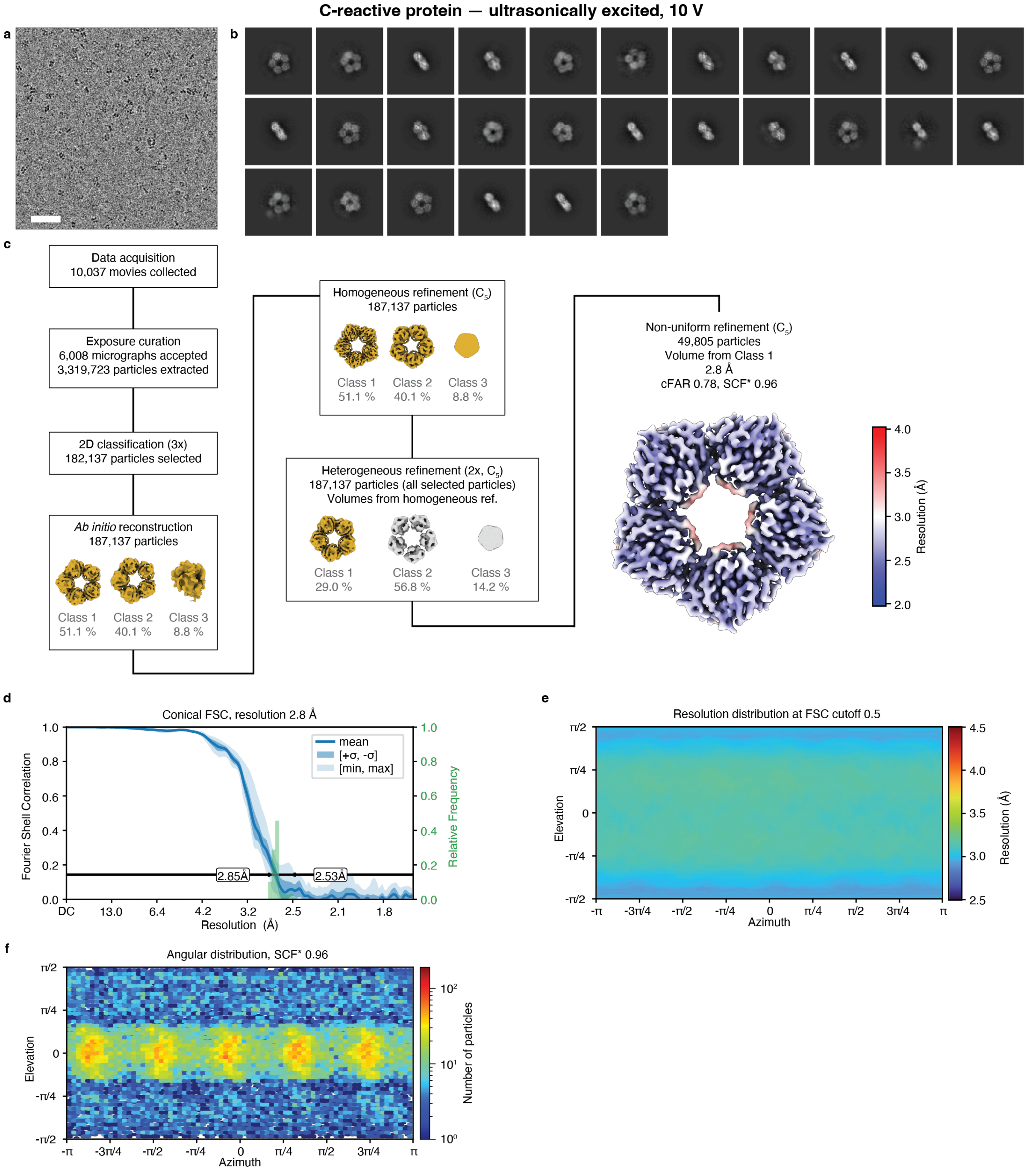


**Figure S10 | Processing workflow for the ultrasonically excited (10 V amplitude) sample of C-reactive protein. a** Representative micrograph. Scale bar, 500 Å. **b** Selected 2D class averages. **c** Data processing workflow in CryoSPARC. Symmetry applied is indicated in the parentheses. The volumes are displayed at a level of 10σ above the mean. The final map is displayed at a level 10σ above the mean, with the local resolution estimation at the FSC cutoff of 0.5 indicated in color. **d** Conical FSC, with the 0.143 threshold indicated by a black line. The mean FSC value is shown as a solid blue line.

Dark blue shading indicates one standard deviation of the FSC curve, while the light blue shading represents the minimum and maximum FSC values. A histogram of the resolution values obtained from the conical FSC is shown in green. **e** Angular distribution of the resolution at FSC cutoff 0.5, generated using the 3DFSC file from Orientation Diagnostics job in CryoSPARC.

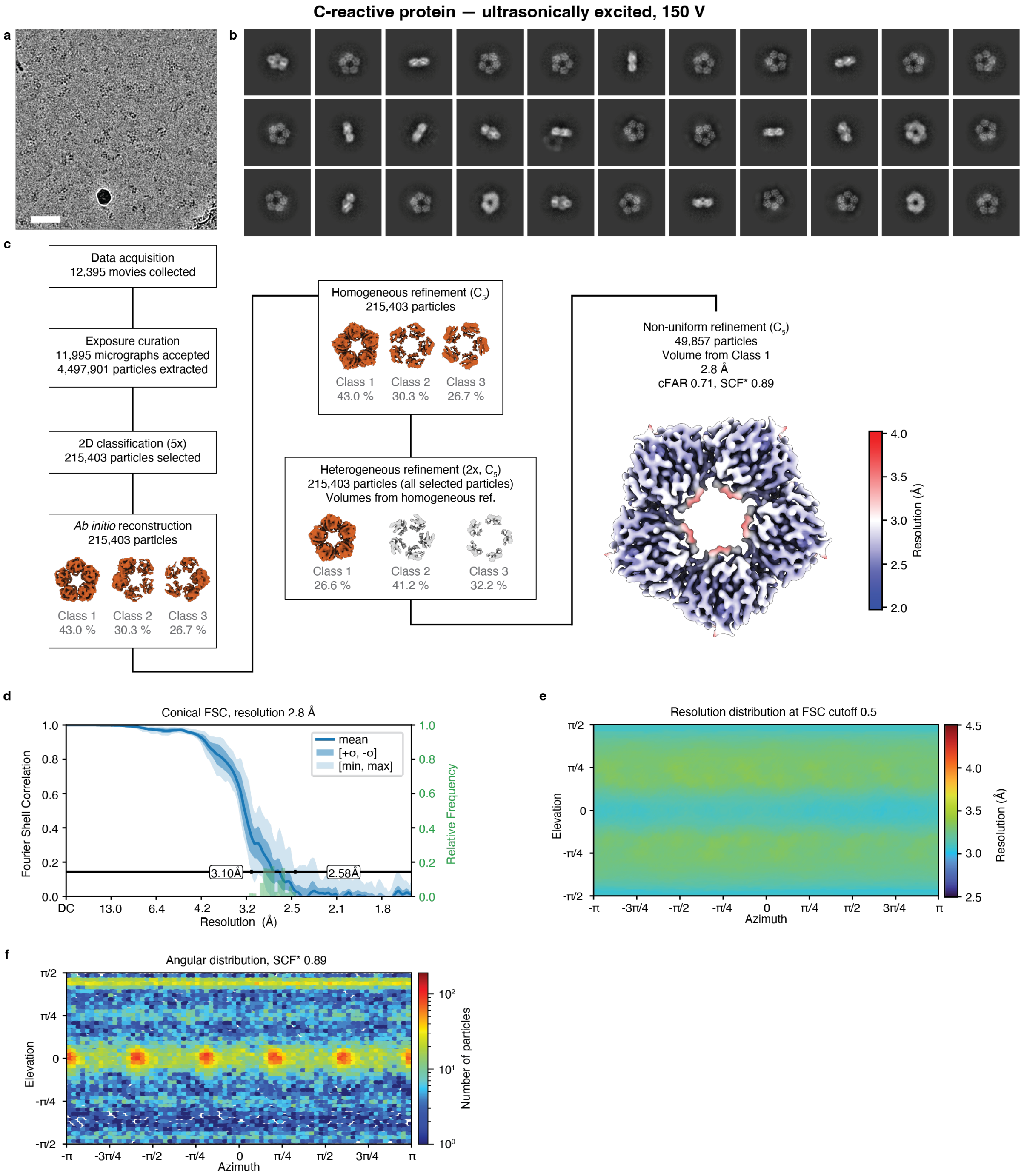


**Figure S11 | Processing workflow for the ultrasonically excited (150 V amplitude) sample of C-reactive protein. a** Representative micrograph. Scale bar, 500 Å. **b** Selected 2D class averages. **c** Data processing workflow in CryoSPARC. Symmetry applied is indicated in the parentheses. The volumes are displayed at a level of 10σ above the mean. The final map is displayed at a level 10σ above the mean, with the local resolution estimation at the FSC cutoff of 0.5 indicated in color. **d** Conical FSC, with the 0.143 threshold indicated by a black line. The mean FSC value is shown as a solid blue line.

Dark blue shading indicates one standard deviation of the FSC curve, while the light blue shading represents the minimum and maximum FSC values. A histogram of the resolution values obtained from the conical FSC is shown in green. **e** Angular distribution of the resolution at FSC cutoff 0.5, generated using the 3DFSC file from Orientation Diagnostics job in CryoSPARC.

**7 | Data processing and analysis – 50S ribosomal subunit**

High-resolution micrographs of all samples of 50S ribosomal subunit (conventional and ultrasonically excited) were collected at the Dubochet Center for Imaging in Lausanne, Switzerland, using a Thermo Fisher Titan Krios G4i transmission electron microscope equipped with a Falcon IV direct electron detector. Single particle reconstructions were performed using CryoSPARC v5.0.6[1] with the corresponding workflows illustrated in Figs. S12–S14. The data acquisition parameters and processing statistics are summarized in Supplementary table 3. Visualization of the density maps was performed using Chimera X[2,3].

Micrographs were first subjected to patch motion correction and patch CTF estimation. Micrographs with estimated CTF fit worse than 5 Å were rejected. Particles from conventional and ultrasonically excited samples were picked using a blob picker (diameter 150 – 400 Å), extracted with a box size of 784 px and Fourier-cropped to 392 px.

All datasets were processed using the identical protocol. First, the extracted particles were subjected to two rounds of 2D classification, followed by an ab initio reconstruction using 3 classes ($C_1$ symmetry). The best of these classes was homogeneously refined ($C_1$ symmetry). The obtained volume was used together with 3 decoy classes and all extracted particles as an input into 3 rounds of heterogeneous refinement ($C_1$ symmetry). Finally, 100,000 particles belonging to the best class were picked randomly, re-extracted using a box size of 588 px and homogeneously refined ($C_1$ symmetry), producing the final volumes shown in Fig. S12 (conventional) and Figs. S13 and S14 (ultrasonically excited with 10 V and 150 V amplitude, respectively).

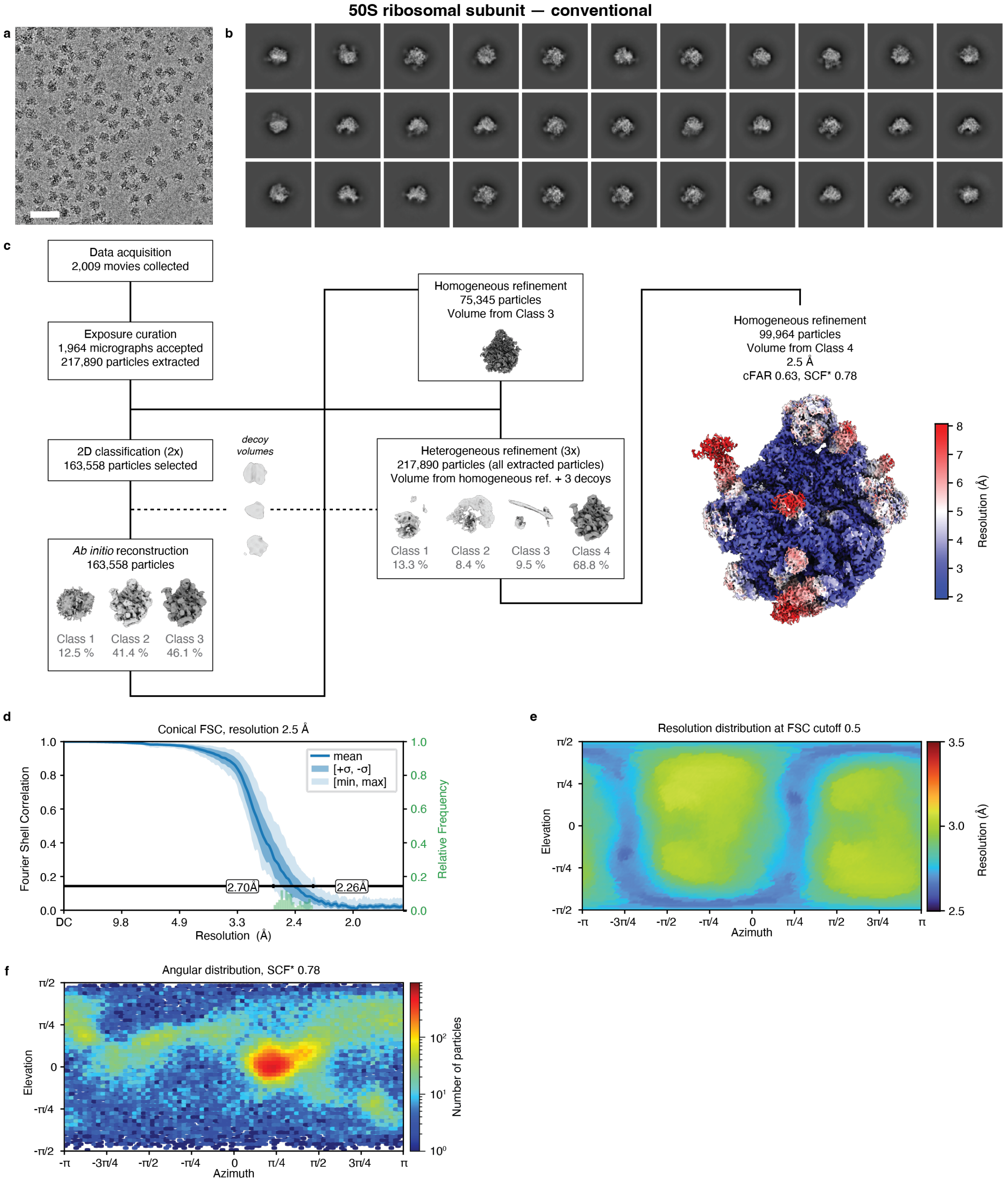


**Figure S12 | Processing workflow for the conventional sample of the 50S ribosomal subunit.** **a** Representative micrograph. Scale bar, 500 Å. **b** Selected 2D class averages. **c** Data processing workflow in CryoSPARC. All reconstructions are performed with $C_1$ symmetry. The volumes are displayed at a level of 5σ above the mean. The final map is displayed at a level 2.5σ above the mean, with the local resolution estimation at the FSC cutoff of 0.5 indicated in color. **d** Conical FSC, with the 0.143 threshold indicated by a black line. The mean FSC value is shown as a solid blue line. Dark blue shading indicates one standard deviation of the FSC curve, while the light blue shading represents the minimum and maximum FSC values. A histogram of the resolution values obtained from the conical FSC is shown in green. **e** Angular distribution of the resolution at FSC cutoff 0.5, generated using the 3DFSC file from Orientation Diagnostics job in CryoSPARC. **f** Angular distribution of the final reconstruction.

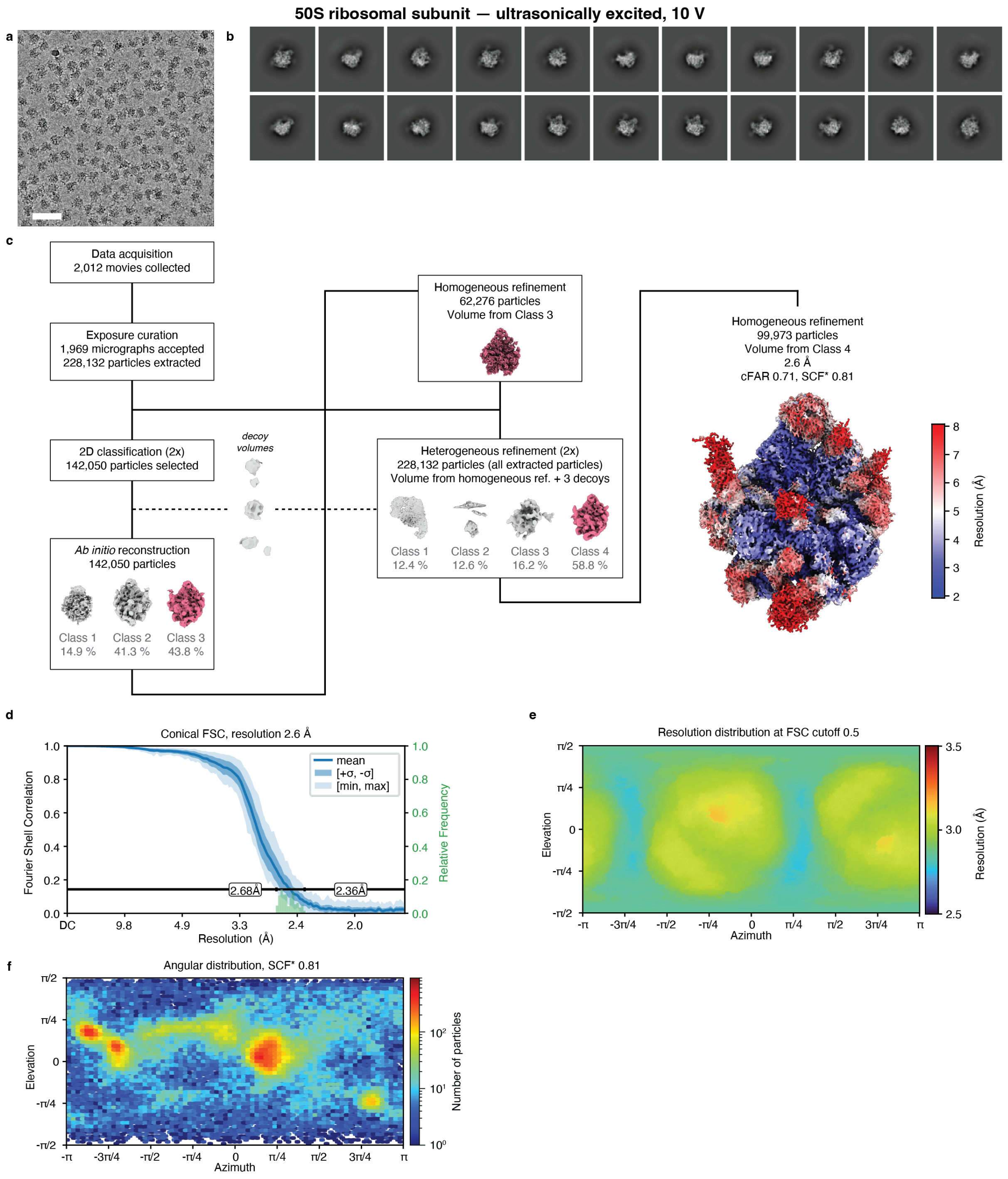


**Figure S13 | Processing workflow for the ultrasonically excited (10 V amplitude) sample of the 50S ribosomal subunit.** **a** Representative micrograph. Scale bar, 500 Å. **b** Selected 2D class averages. **c** Data processing workflow in CryoSPARC. All reconstructions are performed with $C_1$ symmetry. The volumes are displayed at a level of 5σ above the mean. The final map is displayed at a level 2.5σ above the mean, with the local resolution estimation at the FSC cutoff of 0.5 indicated in color. **d** Conical FSC, with the 0.143 threshold indicated by a black line. The mean FSC value is shown as a solid blue line. Dark blue shading indicates one standard deviation of the FSC curve, while the light blue shading represents the minimum and maximum FSC values. A histogram of the resolution values obtained from the conical FSC is shown in green. **e** Angular distribution of the resolution at FSC cutoff 0.5, generated using the 3DFSC file from Orientation Diagnostics job in CryoSPARC. **f** Angular distribution of the final reconstruction.

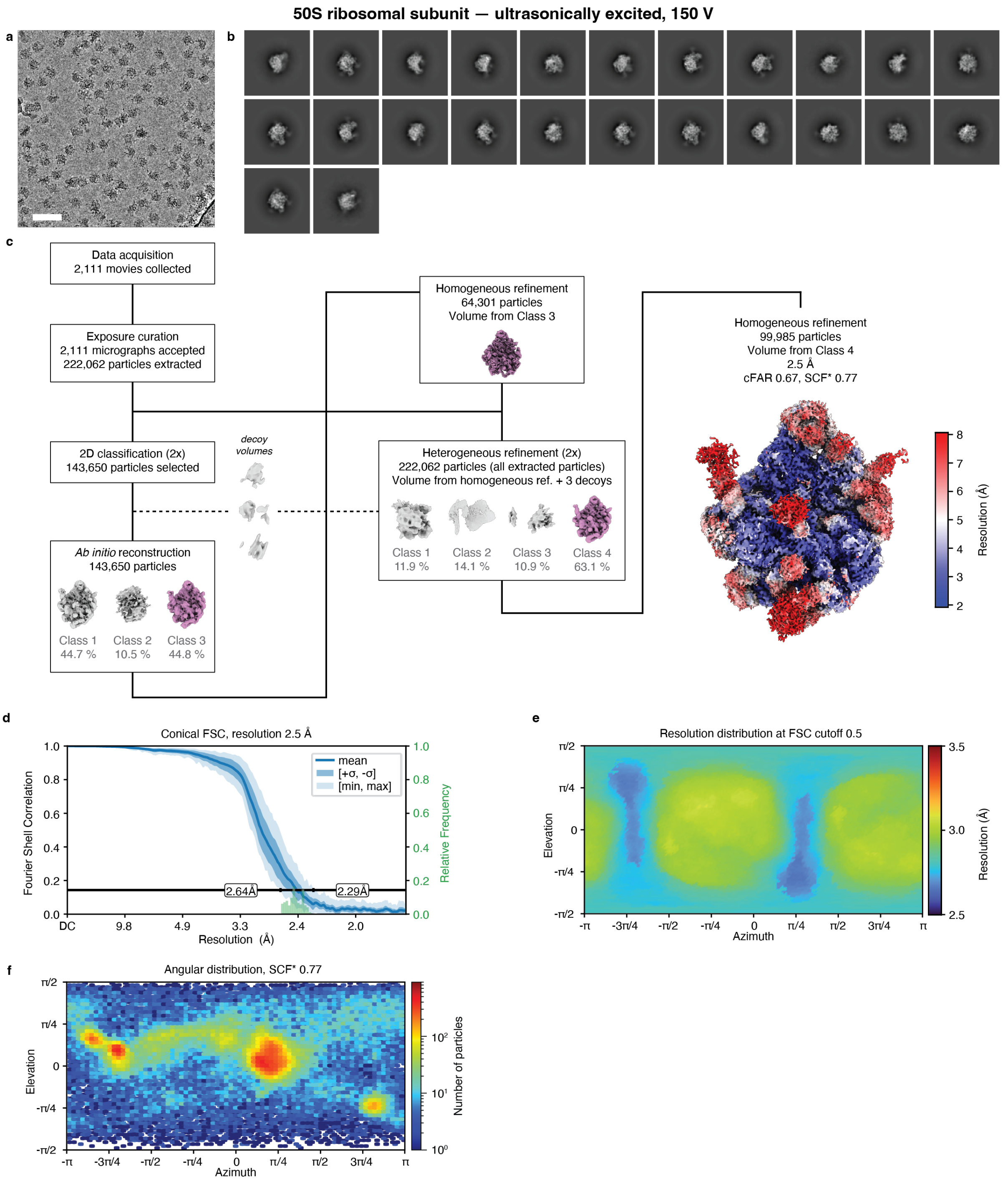


**Figure S14 | Processing workflow for the ultrasonically excited (150 V amplitude) sample of the 50S ribosomal subunit. a** Representative micrograph. Scale bar, 500 Å. **b** Selected 2D class averages. **c** Data processing workflow in CryoSPARC. All reconstructions are performed with $C_1$ symmetry. The volumes are displayed at a level of 5σ above the mean. The final map is displayed at a level 2.5σ above the mean, with the local resolution estimation at the FSC cutoff of 0.5 indicated in color. **d** Conical FSC, with the 0.143 threshold indicated by a black line. The mean FSC value is shown as a solid blue line. Dark blue shading indicates one standard deviation of the FSC curve, while the light blue shading represents the minimum and maximum FSC values. A histogram of the resolution values obtained from the conical FSC is shown in green. **e** Angular distribution of the resolution at FSC cutoff 0.5, generated using the 3DFSC file from Orientation Diagnostics job in CryoSPARC. **f** Angular distribution of the final reconstruction.

### 8 | Cryo-EM data collection and processing statistics

**Supplementary table 1:** Data collection parameters and processing statistics for the hemagglutinin samples.

| | Hemagglutinin conventional sample<br><br>(EMD-100126) | Hemagglutinin ultrasonically excited (10 V amplitude)<br><br>(EMD-100127) | Hemagglutinin ultrasonically excited (20 V amplitude)<br><br>(EMD-100133) | Hemagglutinin ultrasonically excited (150 V amplitude)<br><br>(EMD-100134) |
|---|---|---|---|---|
| **Data collection and processing** | | | | |
| Microscope | Titan Krios G4i | Titan Krios G4i | Titan Krios G4i | Titan Krios G4i |
| Camera | Falcon 4i | Falcon 4i | Falcon 4i | Falcon 4i |
| Energy filter | N/A | N/A | N/A | N/A |
| Magnification | 96,000 | 96,000 | 96,000 | 96,000 |
| Voltage (kV) | 300 | 300 | 300 | 300 |
| Electron exposure ($e^{-}/Å^2$) | 50 | 50 | 50 | 50 |
| Defocus range (μm) | -1.0 – -2.4 | -1.0 – -2.4 | -1.0 – -2.4 | -1.0 – -2.4 |
| Pixel size (Å) | 0.830 | 0.830 | 0.830 | 0.830 |
| Image format | EER | EER | EER | EER |
| Symmetry imposed | $C_3$ | $C_3$ | $C_3$ | $C_3$ |
| Initial particle images (no.) | 5,115,332 | 8,902,027 | 6,368,365 | 6,384,392 |
| Final particle images (no.) | 137,150 | 41,316 | 80,202 | 78,171 |
| Map resolution (Å) | 2.9 | 3.2 | 2.6 | 2.5 |
| FSC threshold | 0.143 | 0.143 | 0.143 | 0.143 |
| Map resolution range (Å) | 11.17 – 2.50 | 3.57 – 2.94 | 2.65 – 2.42 | 2.60 – 2.39 |
| Map sharpening *B* factor ($Å^2$) | 80.3 | 85.1 | 83.9 | 77.3 |
| Orientation diagnostics | | | | |
| cFAR | 0.02 | 0.56 | 0.82 | 0.79 |
| SCF* | 0.32 | 0.74 | 0.87 | 0.83 |

**Supplementary table 2:** Data collection parameters and processing statistics for the C-reactive protein samples.

| | CRP<br>conventional sample<br><br>(EMD-100107) | CRP<br>ultrasonically excited<br>(10 V amplitude)<br>(EMD-100124) | CRP<br>ultrasonically excited<br>(150 V amplitude)<br>(EMD-100125) |
|---|---|---|---|
| **Data collection and processing** | | | |
| Microscope | Titan Krios G4i | Titan Krios G4i | Titan Krios G4i |
| Camera | Falcon 4i | Falcon 4i | Falcon 4i |
| Energy filter | N/A | N/A | N/A |
| Magnification | 96,000 | 96,000 | 96,000 |
| Voltage (kV) | 300 | 300 | 300 |
| Electron exposure ($e^-/Å^2$) | 40 | 40 | 40 |
| Defocus range (µm) | -1.0 – -2.4 | -1.0 – -2.4 | -1.0 – -2.4 |
| Pixel size (Å) | 0.830 | 0.830 | 0.830 |
| Image format | EER | EER | EER |
| Symmetry imposed | $C_5$ | $C_5$ | $C_5$ |
| Initial particle images (no.) | 5,305,207 | 3,319,723 | 4,497,901 |
| Final particle images (no.) | 49,703 | 49,805 | 49,857 |
| Map resolution (Å) | 3.3 | 2.8 | 2.8 |
| FSC threshold | 0.143 | 0.143 | 0.143 |
| Map resolution range (Å) | 3.58 – 3.08 | 2.85 – 2.53 | 3.10 – 2.58 |
| Map sharpening *B* factor ($Å^2$) | 119.6 | 90.0 | 97.3 |
| Orientation diagnostics | | | |
| cFAR | 0.62 | 0.78 | 0.71 |
| SCF* | 0.82 | 0.96 | 0.89 |

**Supplementary table 3:** Data collection parameters and processing statistics for the 50S ribosomal subunit samples.

| | 50S<br>conventional sample<br><br>(EMD-100096) | 50S<br>ultrasonically excited<br>(10 V amplitude)<br>(EMD-100097) | 50S<br>ultrasonically excited<br>(150 V amplitude)<br>(EMD-100108) |
|---|---|---|---|
| **Data collection and processing** | | | |
| Microscope | Titan Krios G4i | Titan Krios G4i | Titan Krios G4i |
| Camera | Falcon 4i | Falcon 4i | Falcon 4i |
| Energy filter | N/A | N/A | N/A |
| Magnification | 96,000 | 96,000 | 96,000 |
| Voltage (kV) | 300 | 300 | 300 |
| Electron exposure ($e^{-}/Å^2$) | 40 | 40 | 40 |
| Defocus range (µm) | -0.4 – -1.2 | -0.4 – -1.2 | -0.4 – -1.2 |
| Pixel size (Å) | 0.830 | 0.830 | 0.830 |
| Image format | EER | EER | EER |
| Symmetry imposed | $C_1$ | $C_1$ | $C_1$ |
| Initial particle images (no.) | 217,890 | 228,132 | 222,062 |
| Final particle images (no.) | 99,964 | 99,973 | 99,985 |
| Map resolution (Å) | 2.5 | 2.6 | 2.5 |
| FSC threshold | 0.143 | 0.143 | 0.143 |
| Map resolution range (Å) | 2.70 – 2.26 | 2.68 – 2.36 | 2.64 – 2.29 |
| Map sharpening *B* factor ($Å^2$) | 41.3 | 35.1 | 34.5 |
| Orientation diagnostics | | | |
| cFAR | 0.63 | 0.71 | 0.67 |
| SCF* | 0.78 | 0.81 | 0.77 |

**9 | References**


1. Punjani, A., Rubinstein, J. L., Fleet, D. J. & Brubaker, M. A. cryoSPARC: algorithms for rapid unsupervised cryo-EM structure determination. *Nat Methods* **14**, 290–296 (2017).
2. Goddard, T. D. *et al.* UCSF ChimeraX: Meeting modern challenges in visualization and analysis. *Protein Science* **27**, 14–25 (2018).
3. Pettersen, E. F. *et al.* UCSF ChimeraX : Structure visualization for researchers, educators, and developers. *Protein Science* **30**, 70–82 (2021).
4. Williams, H. M. *et al.* Overcoming Preferred Orientation in Cryo-EM With Ultrasonic Excitation During Vitrification. Preprint at https://doi.org/10.1101/2025.09.14.676144 (2025).
